# Deep Brain Magnetothermal Silencing of Dopaminergic Neurons via Endogenous TREK1 Channels Abolishes Place Preference in Mice

**DOI:** 10.1101/2022.04.12.487994

**Authors:** Junting Liu, Rahul Munshi, Muye He, Sara D. Parker, Arnd Pralle

**Affiliations:** Department of Physics, University at Buffalo, the State University of New York, Buffalo, 14260, New York, USA

## Abstract

Remote neuromodulation techniques have revolutionized our understanding of brain circuits and their role in behavior. The reversible silencing of specific neuronal populations has emerged as a powerful tool to investigate the necessity and sufficiency of these populations in behavioral responses. Here, we apply magnetothermal silencing using endogenous TREK-1 channels to selectively suppress dopaminergic reward in the ventral tegmental area (VTA) to prove that activation of this specific reward circuit is required for place preference in mice. Magnetothermal silencing entails the application of alternating magnetic fields that penetrate tissue, leading to the heating of superparamagnetic nanoparticles at the target cells, without causing any attenuation or adverse effects. The resultant slight, rapid, and reversible elevation in temperature effectively suppresses neuronal firing, without necessitating genetic modification of the neurons. We demonstrate that two-pore potassium channels, TREK-1, are responsible for this thermal neuronal silencing. Using fiber-based optogenetics we measured both the heating and neuronal silencing in the VTA brain of the animals. We show that in a place preference assay, magnetothermal neuronal silencing of the dopaminergic neurons in the VTA is sufficient to abolish the place preference. Notably, TREK1 knock-out mice exhibit immunity to magnetothermal silencing, behaving as if the magnetic field was not applied. These results underscore the critical role of dopaminergic neuronal activity in the VTA for the establishment of place preference and highlight the dependency on functional TREK1 channels in this magnetothermal silencing approach.

**Highlights:**

- TREK1 is a highly efficient, thermally activated neuronal silencer
- First magnetothermal neuronal silencing in behaving mice
- Fiber photometry quantification of local heating and silencing of target neurons in the ventral tegmental area
- Magnetothermal suppression of dopaminergic reward response in the ventral tegmental area is sufficient to abolish place preference

## Introduction

Rodents have an innate tendency of choosing the dark room when presented with a choice between a dark and a bright room [1]. This choice can stem from either an aversion to the bright room, preference for the dark room, or a combination of both. To better understand the underlying mechanisms, the most straightforward approach is to selectively silence the neuronal pathways associated with preference or aversion and assess the resulting impact on behavior. In the past decade, various optogenetic [2–5] and chemogenetic [5–8] silencing tools have been developed to selectively study the roles of specific neuronal populations [9]. The quick onset and short duration of optogenetic neuromodulation are excellent for activation [10–12]. However, neuronal silencing needs to persist for the entire duration of behavior or sensory input which may continue for tens of seconds to minutes. Such prolonged optogenetic silencing exposes the neurons to potentially damaging light intensities for an extended period [13–15]. Chemogenetics, on the other hand, offers a longer-lasting form of neuromodulation, with effects persisting for minutes to hours, and even extending to days and weeks. However, chemogenetic modulation is not sufficiently transient to be precisely timed with specific behaviors. Instead, chemogenetics induce alterations in the brain circuit throughout the entirety of an experiment and beyond [16].

Our research, along with other studies, has demonstrated that magnetothermal-genetic activation offers neuromodulation capabilities that are optimally suited for precise and timely silencing [17,18], [19] This approach brings several advantages, including its tetherless nature, the ease of targeting large or multiple brain regions, and the ability to define the stimulation location within the experimental arena by manipulating the position of the external magnetic field. Recently, we demonstrated magnetothermal silencing of neurons in culture by overexpressing the chloride channel ANO1/TEM16A [20]. In this work, we demonstrate the first magnetothermal neuronal silencing in behaving mice utilizing the endogenously expressed potassium leak channel TREK1 [21,22].

The two-pore (K2P) channel, TREK1, a TWIK − related potassium channel [23], is not only mechanosensitive [24], [25] but also thermosensitive [26]. Prior work showed that overexpressing a constitutively active TREK1 channel silences hyperactive neurons and ameliorates epileptic seizures [27]. Therefore, the TREK1 channel should be well-suited as a thermal silencer for neuronal activity [26]. TREK1 is expressed in the central nervous system (CNS), cortex, dorsal root ganglia (DRG), and hippocampus. Generally, high expression levels are found throughout the midbrain, although the expression is moderate in the ventral tegmental area (VTA) [28,29]. TREK1 channels are thought to play a role in tuning the activation threshold of neurons involved in depression [30], glutamate release in astrocytes [31], and responsiveness to volatile anesthetics [32].

In this study, we employed TREK1-based magnetothermal neuronal silencing to investigate whether silencing dopaminergic reward alone is sufficient to eliminate the preference in a decision-making assay. Conditional place preference between a dark and a well-lit room was used as a simple decision paradigm. Magnetothermal neuromodulation proves to be exceptionally well-suited for this assay since the magnetic field can be selectively applied to only one of the rooms, establishing a direct correlation between stimulation and the animal’s location. Furthermore, the duration of the animal’s stay in a particular location aligns favorably with the temporal resolution of magnetothermal neuronal silencing.

## Results

### Entry into the dark room in a light/dark chamber activates dopaminergic neurons in the Ventral Tegmental Area

We hypothesized that the innate preference of rodents for the dark chamber in a light/dark test may be attributed more to a positive reward response upon entering the dark area rather than a mere aversion to the brightly illuminated regions [33]. To determine whether a positive reward response occurs upon entering the dark area, we quantified the activity of dopaminergic neurons (DA) in the ventral tegmental area (VTA) of mice. Initially, we examined the neuronal activity of mice in two different settings: an evenly brightly lit arena and an arena divided into a dark and a light room (Fig. 1 A - D). Subsequent c-Fos staining revealed a significantly higher number of active neurons in the VTA of mice in the dark/light chamber compared to those in the light-only chamber, while the total number of cells and dopaminergic neurons remained unchanged (DAPI and TH stains in Fig. 1 D, and Suppl. Fig. S1 A-D). The majority (65%) of neurons in the VTA brain area are dopaminergic [34]. Detailed analysis revealed that the vast majority of the C-fos positive neurons were dopaminergic, with an 86% overlap with TH-positive neurons. Additionally, the activity in GABA-positive neurons was unchanged (Suppl. Fig. S1 E,F). This data demonstrates that having access to a dark room during the experiment leaves the imprint of a positive reward in the animal’s VTA.

**Figure 1:**
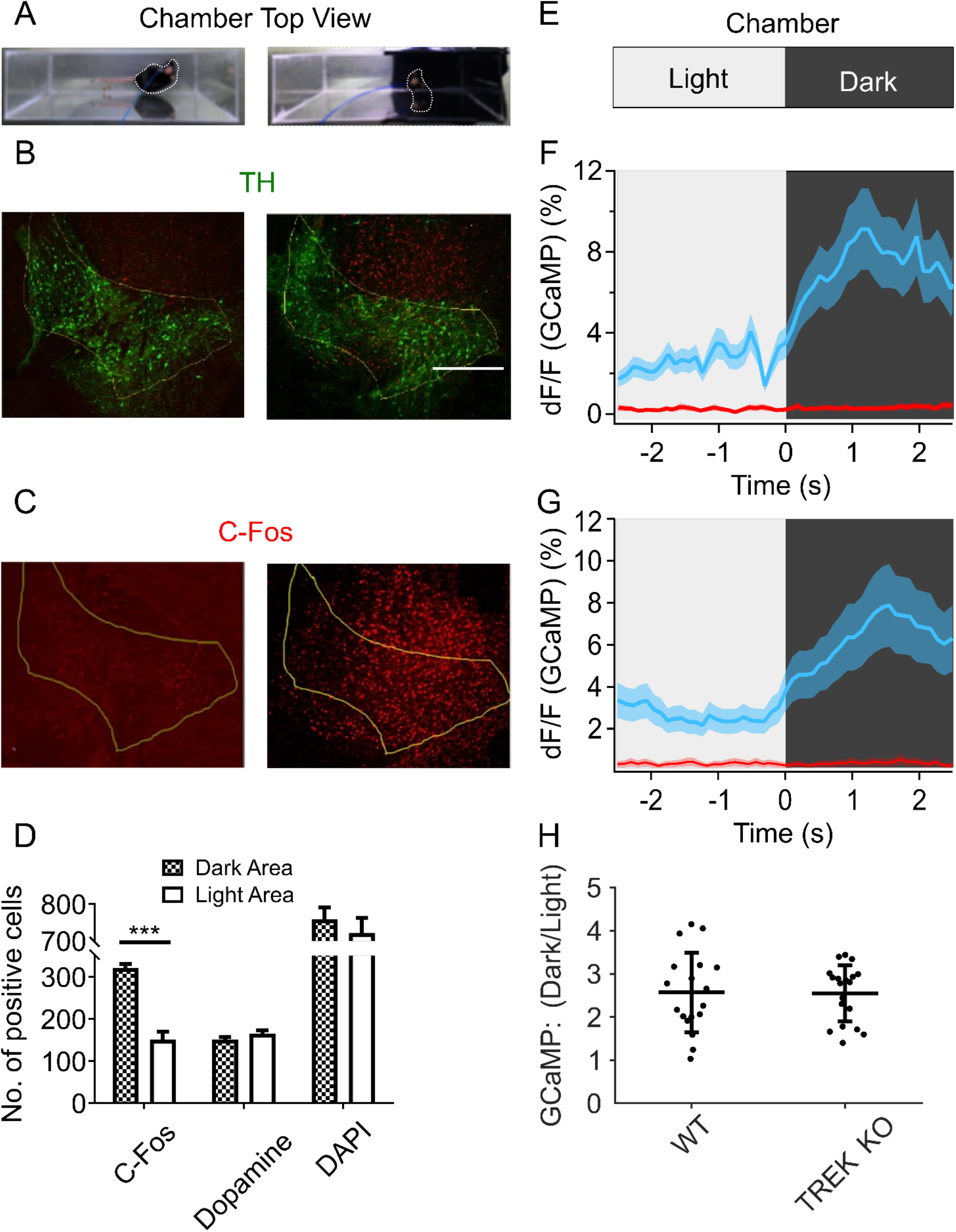
Entering the dark in a dark/bright assay activates VTA DA neurons. **(A)** Light-dark behavior chamber setup schematic, place-preference assay for intrinsic preference of dark space. Half chamber is dark, and another half is bright chamber. **(B)** Representative TH/C-Fos staining in the lateral VTA profile for mouse dwelling in dark and bright area, scale bar 500 μm. **(C)** C-Fos staining for active neuron during assay: the animal remained in either light or dark arena for 20 min. **(D)** The quantification of DAPI staining of cell nuclear in both dark and bright areas, and the quantification of dopamine neurons (TH staining) number both dark and bright areas. Based on the similar quantification of the nuclear and dopamine cell number in the ROI, The significant quantitative analysis of the different numbers of positive cell number in the same lateral ROI area for a relatively long duration of the mouse the dark and bright area. (***P<0.001, n = 15 from 3 mice/group). **(E)** Animal with implanted fiber in light-dark chamber. **(F)** Per-event plot of averaged calcium signals recording via fiber photometry during bright to the dark area for instant state change (2.5s bright and 2.5s dark area), solid line and the shaded region are the mean ± s.e.m. (n=3 bouts from 2 WT mice). Wild-type animals. (**G)** Same as (F) but for TREK1 knock out animals. **(G)** Summary of dark/light activity from WT and TREK KO animals. Bar plots show the significant change during the state change from dark to bright or reversible bright to dark process, mean ± s.e.m. (***P<0.001, n = 3 from 2 mice/group).

To explore the temporal connection between entering a particular room and the initiation of reward, we employed fiber-photometry. The mice were genetically engineered to express GCaMP6s specifically in the ventral tegmental area (VTA), and an optical fiber was surgically implanted through the skull to accurately measure calcium signals from the VTA. This experimental setup allowed us to record neuronal population activity in real-time as the animals freely roamed and explored an environment that was divided equally into a dark room and a light room (Fig. 1 E). By analyzing the calcium signals, we observed that upon entering the dark room, there was a rapid increase in neuronal activity within the VTA. This increase reached its peak approximately one second after entering the dark room. The kinetics and dynamics of this heightened activity were consistent in both wild-type (WT) animals and TREK1 knockout (KO) animals, suggesting that the observed response is independent of the TREK1 channel function (Fig. 1 F). Further quantification revealed an approximately 2.5-fold increase in neuronal activity in both animal groups (Fig. 1 G, activity during 0.35s to 2.15s versus −2.15s to −0.35s). During the movement of the animals in the opposite direction, from the dark room to the light room, the photometry recordings demonstrated a gradual reduction in neuronal activity in the VTA, returning to the baseline level (Suppl. Fig. S2).

Together, these data support the hypothesis that a positive reward response upon entering the dark room plays a significant role in the preference of rodents for the dark over the light room. To establish a causal relationship between this reward signal and behavior, further investigation involving the suppression of dopaminergic neuronal activation is needed. This can be best achieved by inhibiting the activities of neurons responsible for the positive reward response.

### The TREK1 channel is a fast and strong thermally activated neuronal silencer

To effectively inhibit the activity of dopaminergic neurons in the ventral tegmental area (VTA), we developed a technique known as magnetothermal neuronal silencing. Our previous work demonstrated successful magnetothermal silencing using the chloride channel TMEM16A in vitro [20]. However, we opted against using TMEM16A in vivo due to concerns regarding the potentially detrimental effects of additional chloride influx on neuronal health, as well as the challenges associated with viral packaging due to the size of TMEM16A. Instead, we utilized the potassium leak, two-pore channel TREK1 Instead, we turned our attention to the two-pore channel TREK1, a potassium leak channel. Expression of a constitutively active TREK1 channel in specific regions of the central nervous system (CNS) has been shown to effectively silence hyperactive neurons and mitigate epileptic seizures [27]. As TREK1 is mildly thermosensitive [26], we hypothesized that combining TREK1 with remotely heating magnetic nanoparticles would create a potent and safe magnetothermal neuronal silencer.

To determine whether thermally activated TREK1 would hyperpolarize neurons sufficiently to suppress their activity, we over-expressed TREK1 (TREK1 OE) in cultured hippocampal neurons. The culture was subjected to temperature ramps from 34°C to 42°C. The neuronal spiking measured via GCaMP6f fluorescence subsided immediately upon initiation of the temperature ramp (Fig. 2 A, rate of temperature rise is 0.15 °C/s). Notably, this silencing effect was observed even at bath temperatures below physiological levels. During the cooling phase, the firing of the cells resumed around physiological temperatures. These findings demonstrate that TREK1 OE can sufficiently hyperpolarize neurons and effectively suppress their activity in response to heating.

**Figure 2:**
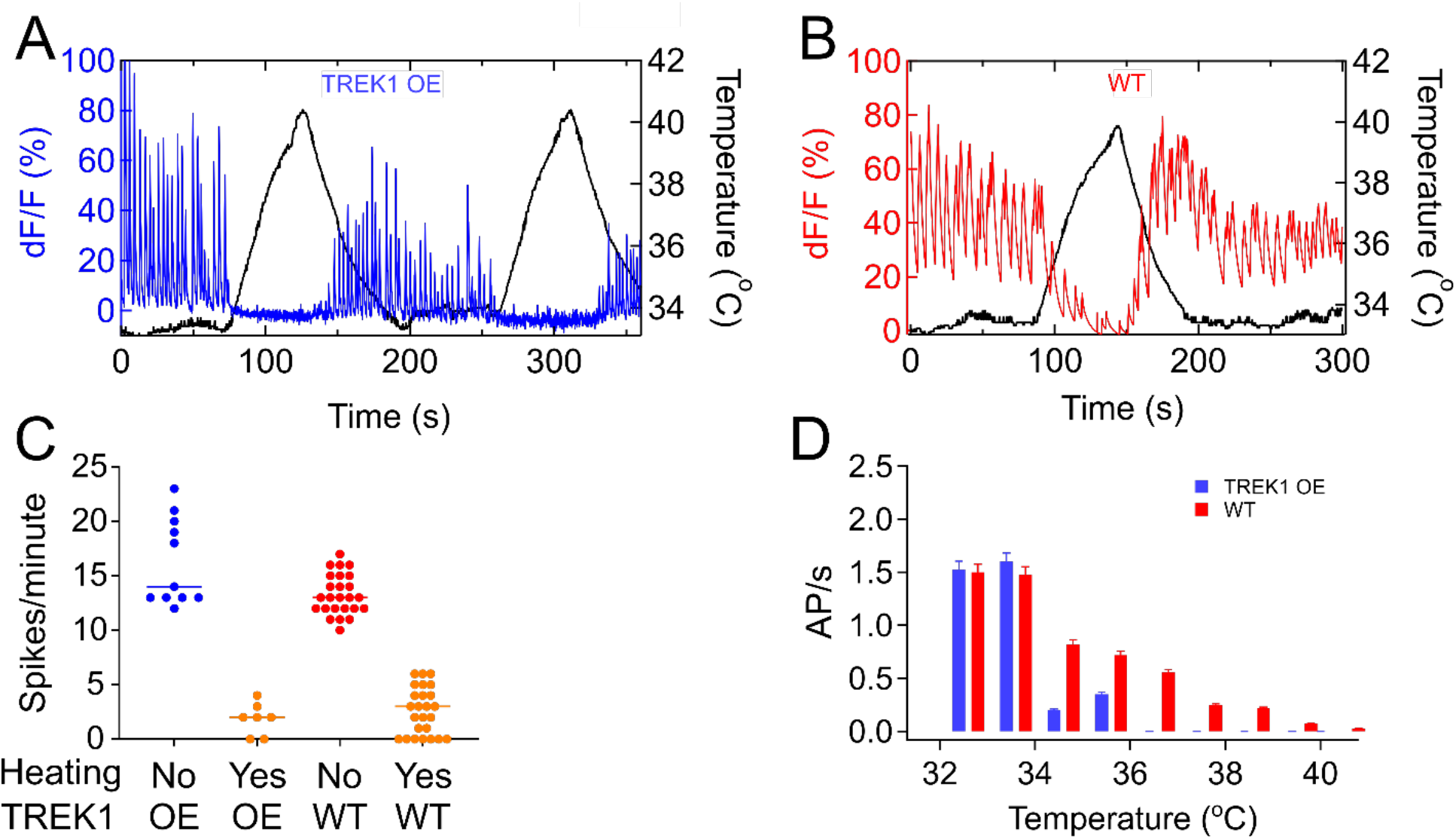
The TREK1 channel is a fast and potent thermally activated neuronal silencer. **(A)** Normalized GCaMP6f fluorescence signal showing silencing of TREK-1 overexpressing (OE) rat hippocampal neurons with positive temperature transients (0.15 °C/s). **(B)** Normalized GCaMP6f fluorescence signal showing silencing of wild-type (WT) rat hippocampal neurons with bath temperature ramp (0.15 °C/s). **(C)** TREK1 overexpressing neurons’ activity as spikes per firing spike numbers per minute without heating (blue, n=11) and spikes per minute during a heating period (heating rate: 0.15 °C/s, n=5). WT neuron’s firing frequency as spikes per minute without heating (red, n=25) and WT cell’s spikes per minute during the heating (heating rate: 0.15 °C/s, orange, n=24) (Solid thin mid-line and the top-down thick lines are the mean ± SEM,. *** p<0.001, two-way ANOVA between without heating and with heating). **(D)** Action potential (AP) rate of WT (red) and TREK1 overexpressing (blue) cultured neurons temperature at difference absolute temperatures (0.15 °C/s heating rate).

In several regions of the central nervous system, TREK1 is expressed endogenously. High expression levels occur throughout the midbrain, with moderate expression in the VTA, hippocampus, cortex, and dorsal root ganglia [21,22,35,36]. Figure 2 B shows that endogenously expressed TREK1 in rat hippocampal neuronal culture suffices as a thermal silencer. The thermal silencing response of WT neurons was slower and slightly weaker than in TREK1 OE neurons. The pooled data of several cultures showed a significant reduction in the number of calcium spikes in both wild-type and TREK1 OE hippocampal neurons when subjected to a positive temperature ramp, with a 5- to 6-fold decrease (Fig. 2 C). Interestingly, in TREK1-OE cells, the thermal silencing effect was efficient even at temperatures below the physiological range, but only in the presence of a fast temperature ramp (Fig. 2 D).

In TREK1-OE cultures the neuronal firing ceases as soon as the temperature ramp begins. The delay between the start of heating and the onset of silencing was 0.9 seconds (n = 13) for TREK1 OE cells and ∼3 seconds (n = 7) for the WT neurons. At a temperature ramping rate was 0.15 °C /s, the neuronal activity ceased before the temperature had risen by 0.15 °C and 0.5 °C in TREK1 overexpressing and WT cells, respectively. This response is significantly faster than magnetothermal activation using TRPV1 [37] and sufficient to establish a strong temporal correlation with behavioral changes.

### *In vivo* magnetothermal heating

In vivo magnetothermal neuromodulation requires precise and controlled heating of specific cell types in the brain. Low radio-frequency alternating magnetic fields (AMFs) enable penetration into the body without substantial attenuation and thus allowing signal delivery into the deep brain without a major loss of penetration depth [38]. The AMFs can then produce produced deep-tissue heating as a function of the magnetic field parameters and the MNPs magnetization [39] (Fig. 3 A, B). The heating modulates the activities of the cells which can then be monitored through the changes in signals of calcium indicators, genetically encoded in the cells. Furthermore, the resulting behavioral changes induced by this modulation can be independently monitored.

**Figure 3:**
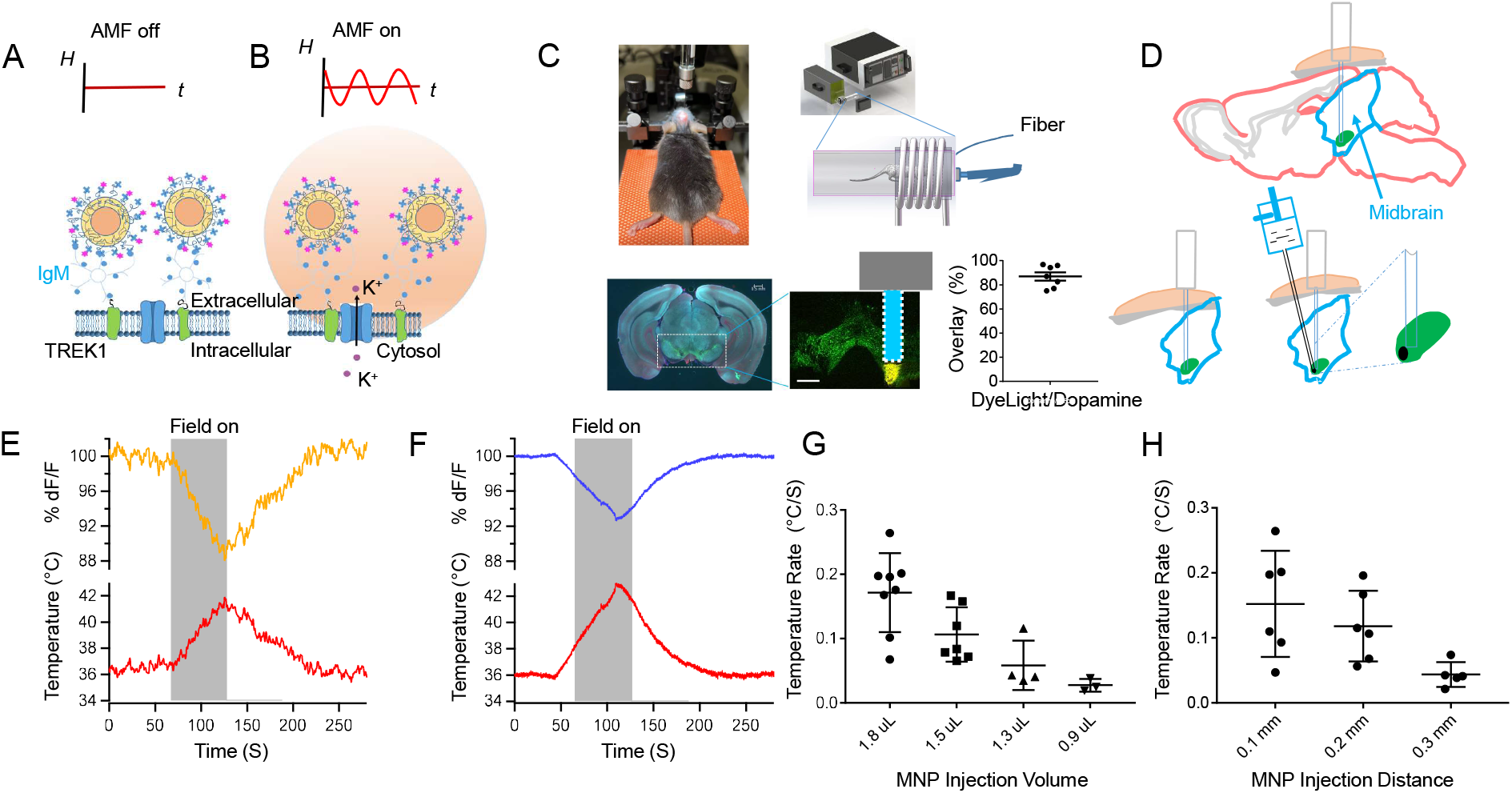
*in vivo* Magneto-thermal heating with Nanoparticles. (A) Alternating Magnetic Field (AMF) stimulation “Field off” of TREK1 without MNP heating is no significant K^+^ flowing out the cell, correspondingly, no change of neuron’s activity. (B) The membrane was heated with MNPs upon applying AMF stimulation “Field On” of TREK1, K^+^ flowing out the cell, the neuronal activity decreased, and thermal neuro-silencing happened with TREK1 ion channel heating. (C) Top row left: Stereotactic injection of GCaMP<u>7f</u> Virus. Top row right: Schematic diagram of the relative position of the magnetic induction coil and the anesthetized mouse, a custom build, water-cooled coil driven by a power supply, the coil has five turns. The mean magnetic flux density along the mouse is 22.4 kA/m at 412.5 kHz for all the heating experiments. The head of the mouse is approximately at the center of the coil. Bottom row left: VTA profile distribution in the whole brain slice of the coronal plane. Bottom row middle: partially enlarged view of the entire VTA area in the white rectangle of left plot, dashed white rectangles indicate fiber location, the yellow area indicates MNPs distribution, scale bar indicates 400 μm. Bottom row right: The percentage of DyLight 550 (antibody bound the nanoparticles) divides the total number of the dopamine neurons. (D) Top: The implanted fiber relative location targets to the bottom part of the VTA brain region. Bottom: Bulky injection of MNPs with angle direction beside the implanted fiber and the partially enlarged view of the injected MNPs in the VTA region. (E) Top curve: direct measurement of fluorescence change (orange signal) for the DyLight 550 signal bound with an antibody with the nanoparticles in VTA for *in vivo* mouse. Bottom curve: the change in absolute temperature (red signal) corresponding to the change in fluorescence with the calibration factor 1/2.02 of fluorescence change. These temperature measurements are from nanoparticles bound with antibodies. (F) Top curve: direct measurement of fluorescence change (blue signal) for the Gcamp7f signal of dopamine neurons in VTA for *in vivo* mice. Bottom curve: the change in absolute temperature (red signal) corresponding to the change in fluorescence with the calibration factor 1/1.04 of fluorescence change. These temperature measurements are direct from neurons with the bulky injection of MNPs around the neurons. (G) For high density MNP injection, the heating effects with different injection volumes and ∼0.15 mm distance. (H) For the bulky injection of MNPs, the heating effects with different injection distance and volume range 1.3-1.5 μL.

To achieve this, the target cell population was virally infected with gCamp7f (0.5 µl, *AAV9.syn.flex.jGCamp7f.WPRE.SV40*). An optical fiber was immediately implanted over the GCamp7f injection region in the VTA (Fig. 3 C, D). GCamp7f is stably expressed in the cells four weeks after injection. Following the expression time, MNPs were injected along a slightly slanted track relative to the implanted fiber (Figure 3 C, D). These nanoparticles can either remain in a concentrated bolus [18] or bind to the membrane of target neurons if coated with antibodies [37] (Fig. 3 A). When an alternating magnetic field is applied, the nanoparticles generate heat through hysteresis losses, which dissipates into the surrounding tissue (Fig. 3 B). A fluorescent dye, DyLight 550, was conjugated with the polymer coating of the nanoparticles. As the fluorescence lifetime and brightness of many fluorophores are temperature-sensitive [40,41], changes in fluorescence intensity give the local change in temperature at the location of the target cells. As the temperature rises, the fluorescence intensity of DyLight 550 decreases. Fluorescence photometry, using the implanted fiber allowed for the determination of fluorescence intensity changes DyLight 550, at the site of nanoparticle injection, as well that that of the intracellular eGFP, of gCamp7f in the target cells. Using prior calibrations, it is possible to convert the fluorescence changes into actual temperature changes at the site of the nanoparticle injection (Fig. 3 E), and inside the target cells (Fig. 3 F).

MNPs were delivered in two ways: 1) At a high density (40 – 80 mg/ml) without any antibodies. Heating is a function of the injection volume (Fig.3 G), and the MNPs remained in the intercellular space, effectively heating for 1-2 months. 2) MNPs, functionalized with targeted antibodies, are injected in lower concentration (2 - 5 mg/ml). These MNPs then specifically bind to the membrane proteins of the target neurons, which is particularly important, as it enhances the specificity of stimulation, confining the effect to the targeted cells only, without any spillover to the neighboring tissues. The downside of this type of injection is that the nanoparticles are endocytosed rapidly compared to method 1.

For the *in vivo* mouse local heating experiments, the anesthetized/awake mouse was placed in a chamber flanked by the magnetic field generating coil (Fig. 3 C). The chamber was held at a constant temperature of 36 °C. This was achieved by a combination of a water-cooling system and efficient insulation. One minute of AMF application raised the temperature by 6.5 °C, from 36.0 °C to 42.5 °C (Fig. 3 E), in the case of MNPs injected in high density. A similar temperature rises of 5.6°C (from 36.2 °C to 41.8 °C) was observed at the MNP surface, in case of the cell membrane targeted injection. We also repeated the cell-membrane targeted heating experiments in other regions of the brain, namely the motor cortex as well, where a ∼5.5°C rise in temperature was seen.

### Acute *in vivo* magnetothermal silencing of VTA Reward Response

Next, we sought to confirm if magnetothermal heating was sufficient to suppress the activation of the DA neurons in the VTA from a positive reward. The activation of the DA neurons of the VTA was achieved either by an ethanol reward or by a social reward. For both conditions, the DA neurons of the VTA were made to express GCaMP6f (AAV5.Syn.GCamp6f.WPRE.SV40), and the fluorescence signal from behaving mice was photometrically recorded using the implanted fiber (Fig. 3 D).

For the ethanol reward experiment, the mice were placed in a chamber flanked by the AMF generating coil and following a brief habituation, the ethanol reward was provided. Continuous photometric recording of the GCaMP6f signal was done throughout the experiment. Upon providing the ethanol reward, there was a dramatic increase in GCaMP6f signal. Upon an immediate application of AMF, a sharp decline in GCaMP6f was observed (Fig. 4 A). The signal reverted to baseline, some time after turning the AMF off (Fig. 4 A, B).

**Figure 4:**
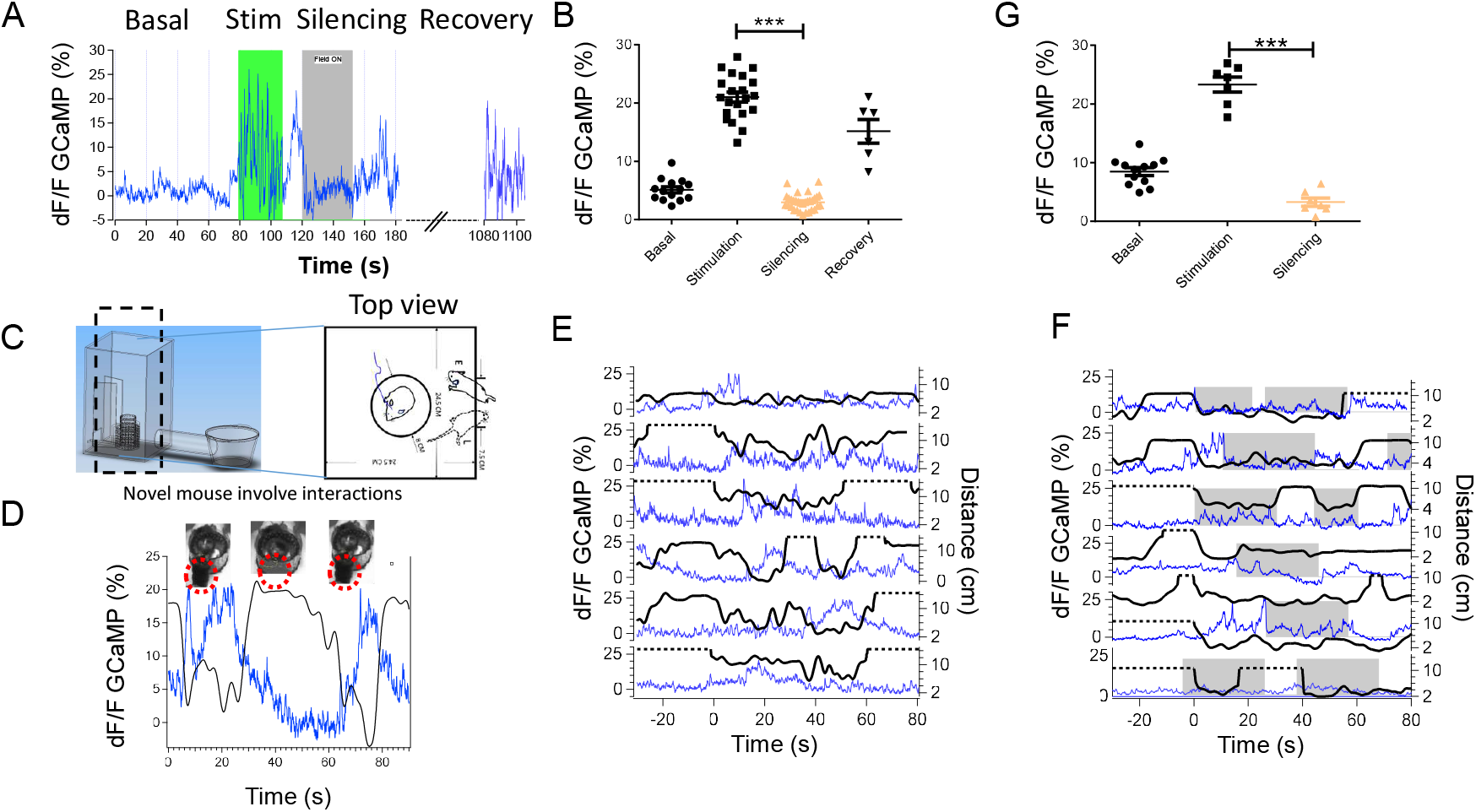
In-vivo magnetothermal silencing of DA neurons in the VTA suppresses reward response. (A) Representative GCaMP6s fluorescence intensity plot from the VTA DA neurons of a mouse. The initial segment of the trace represents the baseline activity of the VTA DA neuronal population, followed by ethanol stimulation (green background), and later followed by magnetothermal silencing (grey) (AMF = 25 kA/m, 425 kHz). The recovery phase following magnetothermal silencing can also be seen in the trace. (B) Scatter plot of different neuronal activity states: basal, ethanol stimulation, magnetothermal silencing (15 minutes), and recovery in the acute ethanol stimulation (4 traces from 2 mice). The relative change in GCaMP6f fluorescence was significantly different between ethanol stimulation and magnetothermal silencing. (C) Left: schematic diagram of magnetothermal stimulation social interaction chamber with a satellite launch room, the recorded mouse located in the cylindrical chamber, flanked by the magnetic coil, and another same-gender novel mouse release in the satellite chamber. Right: An example showing the video tracking of the trajectory of the mouse. (D) (TOP) Snapshots showing the subject mouse, with (left and right) and without (center) the stranger mouse interacting. (BOTTOM) The trace of relative distance between the mice is shown below (black), and gCamp6s fluorescence signal from the VTA DA neurons of the subject mouse, recorded simultaneously is shown in blue. (E) Simultaneous GCaMP6f signal (blue) and relative distance (black) traces are shown for 6 trials. (F) Traces, similar to (E) are shown, but with AMF application (AMF = 25 kA/m, 425 kHz), marked by gray background. (G) Scatter plot of different neuronal activity states: basal, ethanol stimulation, magnetothermal. The relative change in gCamp6s fluorescence was significantly different between ethanol stimulation and magnetothermal silencing.

For the social interaction experiments, we designed a social interaction chamber, with a square housing, with a cylindrical chamber at the center. This cylindrical chamber was flanked by the magnetic field-generating coil and the subject mouse was placed inside it. A stranger mouse of the same gender was initially placed in a satellite chamber, connected to the social interaction chamber with a tunnel. The stranger mouse could explore the entire space, excluding the cylindrical chamber where the subject mouse was placed (Fig. 4 C). An overhead video camera recorded the positions of the mice.

The subject mouse expressed GCaMP6f signal in the DA neurons of the VTA and the corresponding fluorescence signal was recorded using fiber photometry. The GCaMP6f signal showed a dramatic increase whenever the stranger mouse came in close proximity to the subject mouse (Fig. 4 D). This rise in signal, associated with social interaction and was highly consistent and repeatable (Fig. 4 E), making social proximity a robust source of positive reward (4 mice, 2 trials each). We sought to suppress this reward magnetothermally. For this, the AMF was turned on as soon as the stranger mouse approached the subject mouse. Remarkably, this caused a suppression in GCaMP6f fluorescence intensity changes, which was previously observed during social interactions (4 mice, 2 trials each) (Fig. 4 F). Thus, magnetothermal silencing effectively and dynamically suppresses positive reward associated with social interactions in mice.

### Abolition of dark place preference by the magnetothermal silencing of the VTA DA neurons

In a prior section, we argued that CPP (conditional place preference) in a light-dark chamber is a result of the positive reward associated with the dark chamber. We then established that positive reward responses can be suppressed by magnetothermal silencing. This led us to question whether the in-vivo magnetothermal reward suppression is mediated by endogenously expressed TREK1 channels, considering their effectiveness in silencing cultured neurons.

We used a modified light/dark CPP chamber [42], consisting of a small dark (reward) compartment (one-third) and a large illuminated (aversion) compartment (two-thirds). The transition of the mouse between the two chambers can be considered a measure of exploratory activity [33]. Following an initial habituation period, an increasing preference for the dark part was observed. The preference for the dark part could be either a result of a reward associated with darkness or an aversion to light. However, following a result presented earlier in this work, we know that entry into the dark chamber results in an increase in the activity of the VTA DA neuronal population, which is associated with reward. Additionally, it has been shown in an earlier work that the mice changed their behavior and spent more time in the “light chamber” after the VTA DA neuronal activity was suppressed with Adiponectin modulation [43]. This leads us to believe that the long dark dwell times in the modified CPP chamber were a result of a positive reward response to the dark. Since the VTA DA neurons are central to the reward pathway [44,45], suppression of activities of this neuronal population should abolish any reward-based preference.

We sought to achieve that suppression with magnetothermal silencing of VTA DA neurons. Each trial involved placing the mice in a light/dark chamber and allowing them to freely move between the light and dark chambers for a duration of 20 minutes. The dark chamber was flanked by an AMF-generating coil (Fig. 5 A). During the trials, the application of the AMF was initiated from the 5th minute and continued until the 15th minute. The assay was conducted twice daily with a 12-hour interval. The first set of experiments was performed before MNP injection, where the AMF application resulted in no heating. The second set of experiments was conducted after MNP injection, where the AMF application led to heating. Since only the dark chamber was flanked by the AMF generating coil, the heating of the VTA DA neurons via the MNPs occurred only when the mice were in the dark chamber. A plot illustrating the dark preference of the mice, as the fraction of time spent in the dark chamber is shown in Fig. 5 D. The MNPs were delivered with antibodies, and the time of injection is set at 0 hours on the x-axis. A comparison of dark preference before (−72 to −24 hours) and after (24 to 72 hours) MNPs were injected is shown in Fig. 5 F and H. The preference significantly reduced after the MNPs were injected (p<0.001, n = 10). This can be attributed to the magnetothermal silencing of the VTA DA neurons when the mice entered the dark chamber (flanked by the AMF coil). The silencing suppresses reward response from the dark, resulting in a loss of preference.

**Figure 5:**
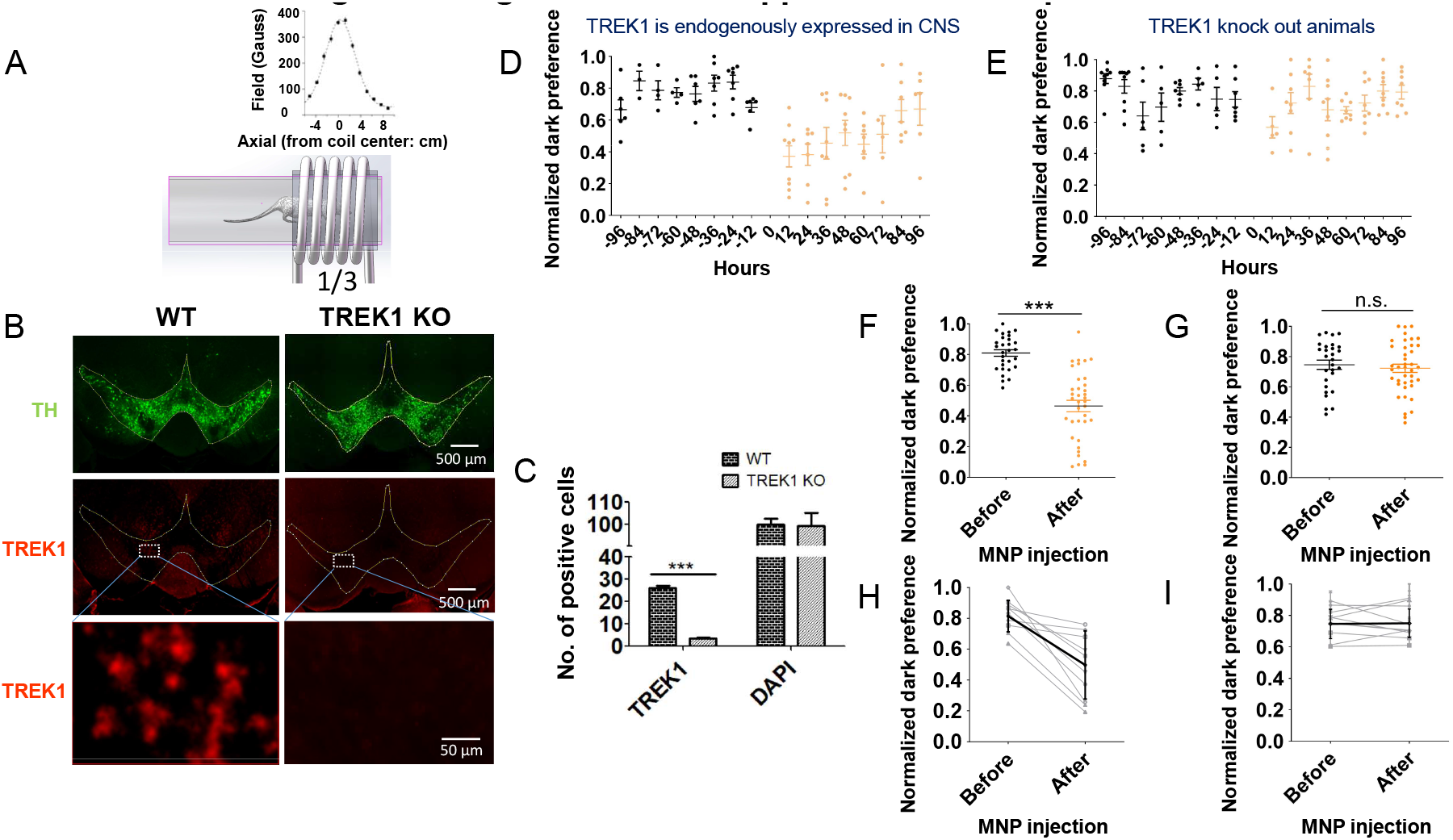
TREK1 mediates Magneto-thermal suppression of dark preference in light/dark CPP assay. **(A)** A schematic diagram showing the modified light/dark CPP assay. The coil covers the dark chamber. The AMF root mean square values at different axial points along the chamber are shown above. **(B)** Micrographs (Left: Wild Type (WT), Right: TREK1 KO) show TH expression (TOP), TREK1 expression (MIDDLE) and a zoom from the dashed boxed area in the middle section. **(C)** Shows the percentage of TREK1 positive cells for WT and TREK1 KO mice. TREK1 KO mice had significantly fewer TREK1 positive cells compared to the WT (p< 0.001). **(D)** The fraction of time the WT mice spend in the dark chamber is given by the normalized dark preference. Data in black were taken at timepoints before MNP injection, while data in orange was taken after MNP injection (n=10). **(E)** The normalized dark preference of TREK1 KO mice. Data in black were taken at timepoints before MNP injection, while data in orange was taken after MNP injection (n=9). **(F)** Comparative scatter plots showing normalized dark preference time before (−72 to −24 hours, black) and after (24 to 72 hours, orange) MNP injection for WT mice (p<0.001). **(G)** Comparative scatter plots showing normalized dark preference time before (−72 to −24 hours, black) and after (24 to 72 hours, orange) MNP injection for TREK1 KO mice (p>0.05). **(H)** Connecting plots comparing before-after pairs of normalized dark preference from each WT mouse. **(I)** Connecting plots comparing before-after pairs of normalized dark preference from each TREK mouse. All error bars represented (mean ± s.e.m)

The period for which magnetothermal silencing is effective depends on the rate of degradation/endocytosis of the injected MNPs. MNPs injected at higher concentration showed greater residence time, that those injected with lower concentration, but with antibodies. However, the antibody-coinjected MNPs are expected to be more precise in heating the desired tissues.

### TREK1 mediates in-vivo magnetothermal suppression of the positive reward response

We hypothesized that the magnetothermal silencing might be mediated by the endogenously expressed TREK1 channels in the VTA. Indeed, there is significant TREK1 expression in the midbrain (Fig. 5 B, C). To establish whether TREK1 is indeed responsible for magnetothermal silencing, a genetic knockout of TREK1 in the brain was performed (Fig. 5 B, Right panels, C). The dark preference of these TREK1 KO mice was similar to the WT mice before the MNP injection. This resulted in the loss of suppression of dark preference in our light/dark conditional place preference (CPP) assay (Fig. 5 E). This is also reflected in the comparison plots Fig. 5 G, I, where the dark preference before (−72 to −24 hours) and after (24 to 72 hours) AMF application was not significantly different (n = 9).

Thus, the magnetothermal silencing of the VTA dopamine response in the dark suppresses the innate preference of the mice for the dark. The loss of preference was truly a loss of preference and not a gain of aversion. The dark preference following magnetothermal suppression was ∼0.5, and during magnetothermal silencing, the duration of time spent in the dark decreased to the one spent in the light. We also verified that this magnetothermal silencing is mediated through the TREK1 ion channels. No change in dark preference was seen in TREK1 KO mice.

## Discussion

In this work, we presented a completely remote and genetic modification-free method of silencing neuronal activity. To demonstrate the efficacy of this technique, we targeted the VTA DA neurons of mice, subjected to various rewarding stimuli, including ethanol administration and social interactions. To showcase the effectiveness of this method, we focused on the dopaminergic signaling in the VTA in mice, which were exposed to different rewarding stimuli, such as ethanol administration and social interactions. By utilizing fiber photometry to capture the signals from genetically encoded calcium indicators, we observed that the calcium signal bursts that typically follow each reward stimulus were rapidly and strongly suppressed using this technique. The inhibition of reward activation holds significant potential for application in animal models related to addiction and gambling disorders.

Furthermore, in order to delve into the long-term consequences of recurrent reward suppression, we investigated whether reward suppression alone suffices to alter the animal’s choice in a dark/light place preference assay. Innately, after acclimatization, mice tend to spend more time in the dark chamber. Our study demonstrated clearly that this behavior arises not from a decline in preference for the “light” chamber, but rather from an increased preference for the dark chamber. We show that this preference stems from an intrinsic reward linked to the mouse’s innate behavior. Leveraging magnetothermal silencing, we succeeded in curtailing this reward-driven activation of VTA DA neurons each time the mouse entered the dark chamber. The mouse’s preference for the dark chamber immediately disappeared, resulting in an equal division of time between the dark and light chambers. This further revealed that the increased dwell time in the dark is a consequence of reward pathway mediated preference, and this preference can be counteracted by inhibiting the activation of the pathway.

Remarkably, the mechanism of silencing does not necessitate the introduction of any proteins through genetic means, as it seems to be primarily facilitated by the endogenous TREK1 ion channels. The endogenous expression of TREK1 proved sufficient to induce potent silencing of excitatory neurons and to induce alterations in behavioral traits. This was demonstrated by the complete insensitivity of the dark place preference among to magnetothermal silencing in TREK1 knockout mice. To understand the implications of TREK1 overexpression, we examined calcium sensitive fluorescence in rat hippocampal cultures. Notably, the activity of neurons with wild-type expression was indeed suppressed upon elevation of the bath temperature. Interestingly, neurons overexpressing TREK1 exhibited an astonishingly rapid and sustained suppression of activity. We observed that this swift silencing hinged on the rate of temperature ramp and not on the absolute temperature itself.

This finding offers the potential to devise a neuronal silencing system that operates through swift heat generation, achievable in exceedingly brief bursts. Consequently, there is no necessity to escalate temperatures to levels that might prove detrimental to neurons. An additional advantage lies in the abundance of TREK1 channels within the brain [29]. In summary, thermal silencing mediated by TREK1 emerges as a highly promising avenue for modulating neuronal activity. When coupled with advancements in the efficiency of magnetic nanoparticle heating [46] and delivery techniques [47], this genetic modification-free magnetothermal silencing emerges as a feasible contender for widespread adoption across various studies that demand a rapid, robust, non-toxic, implant-free, minimally invasive, and genuinely remote approach to neuronal control.

## Materials and Methods

### Animals

Male and female C57BL/6 mice wild-type (WT) (Jackson Laboratory, #000664), DAT: Cre mice (Jackson Laboratory, #006660), and C57BL/6 background TREK-1^−/−^ knockout (KO) mice were obtained from Dr. Andreas Schwingshackl (Ronald Reagan UCLA medical center, UCLA Mattel Children’s hospital), while prague Dawley rats were used for cell culture. All of the animals aged 10-12 weeks were housed four per cage and maintained under controlled temperature (23 ± 3 °C) with a reverse 12-hr light/dark cycle and provided standard rodent chow and water available ad libitum during the duration of the experiments. The animals were acclimatized for seven days before experiments and habituated to handling for at least three days before behavioral testing; for other experiments, the animals would return to the animal room after the experiments on the same day. All experimental procedures for animal care and use were approved by the State University of New York at Buffalo Committee on Animal Care and followed the National Institutes of Health Guide for the Care and Use of Laboratory Animals.

### Cell culture and cellular level thermal silencing

#### Preparation of neurons

For dissociated hippocampal neuron culture preparation, hippocampal neurons were harvested from embryonic day 17 (E17) Sprague Dawley fetuses. Briefly, dissected hippocampal tissue was digested with 50 units of papain (Worthington Biochem) for 6 min at 37 °C, and the digestion was stopped by incubation with ovomucoid trypsin inhibitor (Worthington Biochem) for 4 min at 37 °C. Tissue was gently dissociated with Pasteur pipettes, and dissociated neurons were plated at a density of 20–30K per glass coverslip coated with Matrigel (BD Biosciences), and the neurons were placed in Tyrode’s solution (Sigma Aldrich). After cell adhesion, an additional plating medium was added. Neurons were grown at 37 °C and 5% CO_2_ in a humidified atmosphere. After transfection or nucleofection, the neuronal cultures were placed in the incubator for the next 24-48 hours to express the proteins (GCaMP6f or TREK1). During this time, 1 µM TTX (Tetrodotoxin) was added to the neuronal culture to minimize endogenous activity. The TTX was washed out during imaging and then transfected *in vitro* on days 6-8 (DIV 6–8). The imaging buffer also contained synaptic blockers DLAP-5 25 µM, NBQX 10 µM, Gabazine 20 µM, and the osmolality was adjusted to 310-315 mosmole/L. All measurements on neurons were taken after DIV 16.

#### Targeting nanoparticles for the neurons

Nanoparticles were targeted to the neuronal cell membrane via biotinylated A2B5 antibody (433110, Invitrogen). Neuronal cultures grown on a 12 mm diameter coverslip were loaded in the imaging chamber (ALA scientific), holding 150 µl of the imaging buffer (PH 7.3). Biotinylated antibody (2 µg/ml) was added and incubated for 10 minutes before being washed off by perfusion with HEPES buffer (PH 7.3). Neutravidin conjugated nanoparticles were added at 10 µg/ml, and after 5 minutes, the unbound nanoparticles were washed off. The local temperature rise near the nanoparticles was measured by fluorescence intensity changes of a DyLight 550 fluorophore integrated in the NeutrAvidin coating.

#### Recording magnetothermal neuronal membrane heating

Fluorescence microscopy of membrane decorated nanoparticle bound dye was done with an inverted epifluorescence microscope (Zeiss Axio Observer A1M). The alternating magnetic fields (AMFs) induce eddy currents in the metallic microscope objective and heat the nanoparticles. The fluorescence intensities of molecular fluorophores are sufficiently temperature-dependent to provide a molecular scale thermometer. The membrane-bound nanoparticles heated with an area density-scaling factor were calibrated (1.1×10^-3^ ± 6.7×10^-5^ °C.µm^2^/s).

#### Recording the cultured neuron silencing by bath/magnetothermal heating

We collected the cultured neurons’ spike in dissociated rat hippocampal cultures with significantly decreased magnetothermal temperature ramps and absolute temperature increase. The activity data were recorded on wild-type (WT) rat hippocampal neurons and TREK1 overexpressing rat neurons.

### Stereotactic surgeries

TREK1 knockout (KO) or wild type (WT) mice of 10-12 weeks of age were given isoflurane anesthesia before being laterally injected with premixed 0.5 uL Gcamp6f (*AAV5.Syn.GCamp6f.WPRE.SV40*) and TREK1 (*rAAV9/PAAV-hysn-TREK1*) in the VTA region (VTA, coordinates: −3.3 mm AP, ±0.4 mm ML, −4.5 mm DV, AP -Anteroposterior. ML - Mediolateral. DV - Dorsoventral), as well as the DAT: Cre mice were injected in the VTA with GCamp6s (*AAV5.syn.flex.GCamp6s.WPRE.SV40*). ML and AP coordinates are relative to Bregma, and DV coordinates are relative to the dura mater. Hence injection was slowly performed with a microinjection pump (UMP3, UMC4, World Precision Instruments, Inc., Sarasota, FL, USA) at the speed of 60 nL/min with a 31-gauge needle; after virus injection, the needle was left in place for 10 minutes before slowly retracting it.

### Stereotactic cannula implantation

The experimental mice were individually anesthetized with isoflurane and underwent surgery, were mounted in a stereotaxic device, and their head was secured. Before surgery, Buprenorphine (0.1mg/Kg) was administered routinely through an SC injection. Before drilling the hole at the VTA coordinates, ensure that the skull is carefully leveled via DV coordinate measurements. The implanted fiber needs to be connected to the scientific camera and check the increasing signal during the slow implantation of the fiber down to the VTA area. The fiber can be fixed after finding the maximum signal increase above the top part of the VTA (optical fiber implants were placed above the VTA; usual DV depth is between 4.0-4.3 mm) and then glued with Vetbond adhesive around the surgical skin incision. After drying, apply C&B Metabond where the dental cement will contact the skull, and finally, apply several layers of dental cement to affix the fiber onto the skull. Ensure that the mouse is on a heating pad to maintain its body temperature during surgery; after surgery, return the mouse to its home cage when the mouse is fully awake. Carprofen (5mg/kg) was administered once daily subcutaneously for two days.

### Stereotaxic angle injection for magnetic nanoparticles (MNPs)

The mice underwent stereotaxic bulky volume MNP injection (core-shell: CoFe_2_O_4_-MnFe_2_O_4_ ∼14nm diameter or Magnetite: spherical Fe_3_O_4_∼22nm, 10mg/ml, 1.2uL) in the VTA with the coordinates of (−3.3 mm AP, 0.4 mm ML, 4.8 mm DV) with a 27-gauge needle. We adopted the membrane-binding injection to locally heat the brain cell membrane in live mice, or we stereotaxically injected A_2_B_5_ antibody (1mg/ml, 0.6ul) into the region of interest. The injection was followed by MNP (2mg/ml, 0.6uL) injection into the same spot. The injection speed was a constant 60 nL/min (UMP3, UMC4, World Precision Instruments, Inc., Sarasota, FL, USA). Nanoparticles attached to neurons in the brain showed heating for at least 7 days for the membrane binding method, and the volume injection lasted for 1-2 months. For both of these injection methods, the needle was parallel to the sides of the ferrule of the fiber, and then the drill was mounted into the stereotaxic device. Gently drill a hole through the dental cement with a 16-degree angle with the fiber in the front-rear plane, vertical with the coronal plane (16-degrees from rear to front in the vertical direction). Then, mount the needle drawn with 1.2uL MNPs, slowly go down the predetermined coordinates of (−3.3 mm AP, 0.4 mm ML, 4.8 mm DV) and inject the MNPs slowly into the edge of the VTA region or the Gcamp6 expression area (injection speed, 20nL/min for volume, and 60 nl/min for membrane binding). The needle was retracted slowly and waited for a 10-minute interval after every withdrawal of 0.5mm.

### Fiber Photometry setup

The reconfigurable dichroic mirrors were mounted in removable dichroic filter cage cube holders (CM1-DCH, THORLABS) that enabled two different light sources: one fluorescence excitation 470 nm LED light source (M470F3, THORLABS) to be coupled with fiber optic SMA patch cables (M28L01, Ø400 µm, 0.39 NA, Low OH, THORLABS) and a corresponding collimator. In this configuration, a 470-nm LED filtered with a 470-nm bandpass filter (Thorlabs, FB470-10) was fiber-coupled into the dichroic cube holder and another LED light source 405nm beam (M405F1, THORLABS) with a 405-nm bandpass filter (Thorlabs, FB405-10) was coupled in the same way with another collimator (Thorlabs, F671SMA-405 and AD11F) and a 495nm long pass dichroic mirror (Semrock, FF495-Di02-25 ×36). The excitation LED light produced an excitation spot with a ∼2.5-mm diameter at a working distance of a 20×/0.40-NA objective (Olympus, PLAN 20×), and the objective was fixed on a framed stage. A 400-μm-diameter 0.50-NA fiber (THORLABS, FP400ERT) was used to transport fluorescence excitation. One end of the fiber terminated in an FC/PC multimode connector with a ceramic ferrule (Thorlabs, 30440C1) was fixed on 3-axis MicroBlock compact flexure stages (THORLABS, MBT616D) at the working distance of the objective, and the other end terminated in 1.25-mm-diameter ceramic ferrules (CLFC 440). This ferrule was coupled via a ceramic sleeve (Thorlabs, ADAL1) to a 1.25-mm-diameter ferrule implanted into a mouse head. Fluorescence emission from the fibers was passed through a 520nm bandpass fluorescence emission filter (selected for GCaMP6 recording; Semrock, FF01-520/35-25). The fluorescence image was focused onto the sensor of an sCMOS camera (Andor, Zyla 4.2PLUS) through a tube lens (Thorlabs, AC254-040-A-ML) [48].

The excitation GFP 470nm LED light and the correcting motion-related artifacts present in the 405-nm isosbestic wavelength were applied as a 50ms/50ms switching mode, and switching LED lights were controlled by LED drivers (LEDD1B, THORLABS) through Clampex software (V10.7.03, Molecular Device, LLC) and AXON CNS Digidata 1440A hardware (Molecular Device, LLC). The camera sampling frequency was set at 100 Hz and can evenly collect 5 data points during each switching period and utilize the middle three points for the data processing. The 470 nm and 405 nm signals were separated with the threshold, which was determined from the gap of the 470 nm and 405 nm values.

### Alternating magnetic fields (AMF) of the magnetothermal system setup

Thermal sensitive ion channels respond to the alternating magnetic fields (AMFs) stimulus. A customized magnetothermal system generates the AMFs (30 KA/m) in the low radio frequency regime (400KHz, 10 kHz-1 MHz adjustable), which can penetrate arbitrarily deep regions of the brain and interact minimally with biological tissues. On the other hand, magnetic nanoparticles (MNPs) were injected into the particular brain region in the mouse, where the MNPs were exposed to AMF dissipated heat and thus can trigger heat-sensitive ion channels. We functionalized magnetic nanoparticles for bio-conjugation to the cellular membrane or volume injection for suspension in brain tissue to utilize this. Therefore, the particular neuron population or brain region will achieve the absolute temperature or temperature rate heating during the field.

### *In vivo* measurement of local heating via GCamp7f signal using fiber Photometry

Measurement typically starts 24 hours after MNP injection. The experimental mouse was deeply anesthetized with isoflurane for 40 minutes in the induction box with 2-2.5% isoflurane; the implanted ferrule with a fiber glued in it was then placed the site of nanoparticle decoration. This serves as a port to which the photometry fiber can be connected repeatedly during experiments. For all experiments, the mice were anesthetized and placed within the hyperthermia coil, connected to the primary fiber, which is already connected to the fiber photometry, and then put the mouse into the U shape magnetic coil and make sure the mouse head is in the middle of the coil height. The data collecting period needs the room to be dark or cover the fiber photometry and fiber connecter with a black cloth to avoid light from outside. During the recording, the 470 and 405 nm LED lights to need to be turned on according to the Clampex software protocol, and then start recording the data from the Andor Zyla camera for the required time. Finally, analyze the data and convert it to the temperature increasing curve using the calibration data. A typical measurement session lasted about half an hour, after which the animals quickly recovered and were returned to their home cage.

### Calcium imaging recording of the neuron’s activity through fiber Photometry

Imaging typically started two weeks after fiber implantation surgery and 12-24 hours after MNP injection. Mice were lightly anesthetized (0.7-1% isoflurane), then connect the fiber to the ferrule which was implanted into the mouse brain, gently move the mouse into the U magnetic coil and wait for 40 minutes until the mouse is completely awake (no isoflurane effects), and check the mouse’s regular activity and behavior. Similar to the operation of local heating, the recording of neural activity was performed with the awake mouse, start recording the neural activity, click the Clampex recording button to start the LED light as in the 50/50 ms switching protocol, and then start recording the data using the Andor Zyla camera for 5 minutes or other periods. A typical recording session lasted around half an hour, after which the animals were returned to their home cage.

### Conditioned place preference (CPP) behavior

The animals were acclimatized for 3 hours before the experiments and habituated to handling for at least three days (pre-training, twice per day) before the behavioral test. The CPP behavior chamber was covered in black for one-third of the total length, and the black covered part was set in the magnetic coil and exposed to the AMFs during the field on, and then performed on the mice in the behavior chamber, where the experimental mouse is freely moving in the chamber for 20 min, and applied AMFs during the middle 10 minutes, and the first 5 minutes and the last 5 minutes when the AMF was kept off state. The behavior test was repeated from −96 hours to 0 hours with twelve-hour intervals, and then the mice were given a stereotaxic nanoparticle injection (core-shell: CoFe2O4-MnFe2O4 ∼14nm diameter or Magnetite: spherical Fe3O4∼22nm) in the VTA with the coordinates of (−3.3, 0.4, 4.8 mm) at a 16-degree angle with a 27-gauge needle. Following the nanoparticle injection, mice underwent 0 to +96 hour CPP experiments with the same twelve-hour intervals. These CPP behavior tests were designed for different groups of WT mice (n=10) and the TREK1 KO mice (n=9) with different TREK1 expression levels in the VTA region. The light-dark chamber only light area was illuminated by the flat LED light, and light area activity was adopted with the entire light chamber, which was illuminated with the flat LED light. The mouse remained in space for 20 min/trial, and was trained three times with the trained interval of 12 hours to eliminate the spontaneous exploratory behavior in a novel environment, the mouse mostly spent time in the dark area (sacrificed the mouse to maintain the dark state in 60-90 minutes), and performed subsequent C-Fos staining for the dark training and C-Fos assay. In addition, another entire bright chamber was used for both WT and TREK1 KO mice to test the brightness preference (number of excursions from 1/3 of the bright area of the entire bright area normalized in 5 minutes) compared with the dark preference (number of excursions from 1/3 of the dark covered chamber), and number of excursions from the bright/dark area during the previous field, during field heating, and after the field was normalized per 5 minutes.

### Acute *in vivo* magnetothermal silencing during ethanol activation and social interaction

To achieve dramatic silencing effects from hyperactive neurons after applying magnetothermal silencing for in vivo mouse we activated neuronal activity through ethanol administration or social interactions, and we verified the acute magnetothermal silencing effects after ethanol activation or during social interactions.

All of these mice were used the TREK1 OE in VTA (premixed AAV5.Syn.GCamp6f.WPRE.SV40 and rAAV9/PAAV-hsyn-TREK-1) for both of these acute in vivo magnetothermal silencing experiments.

#### Ethanol activation and magnetothermal silencing

Ethanol stimulates the firing activity of midbrain dopamine (DA) neurons, leading to enhanced dopaminergic transmission in the mesolimbic system through intake from the mouse’s mouth. Ethanol has been shown to enhance the intrinsic pacemaker activity of DA neurons directly; this activation provides us with further magnetothermal silencing, and this silencing can be easily observed through the in vivo fiber photometry recording. TRE1 OE mice were used in these magnetothermal experiments (mixed the TREK1 virus and GCamp6f virus, which were injected into the VTA). For ethanol activation and silencing, the subject mouse was located in the vertical coil, and we started to record the neuronal activity in the VTA through fiber photometry. Secondly, apply ethanol, so it is absorbed into the mouse’s mouth. About 5 minutes later, apply AMF to perform the magnetothermal silencing for this ethanol activation and record all the processes.

#### Social interaction activation and magnetothermal silencing

Similarly, we also activated the neuronal activity by social interactions, and this can realize the social interactions in the chamber around the subject mouse (mixed TREK1 virus and GCamp6f virus were injected into the VTA) with fiber photometry real-time recording and allow the same gender mouse to be released from the satellite chamber. Correspondingly, we recorded magnetothermal silencing during the activated duration of social interactions. Therefore, we can observe dopamine neuron activity changes between the social interactions without magnetothermal stimulation and the magnetothermal silencing with AMF was executed during these social interactions.

### Immunohistochemistry

Mice were overdosed with Ketamine/Xylazine and perfused transcardially with cold PBS, followed by 4% paraformaldehyde (PFA) in PBS. Brains were extracted from the skulls and kept in 4% PFA at 4 °C overnight and then 30% sucrose for 48 hours. Sixty-micrometer coronal slices were taken using a vibrating slicer (Compresstome VF-200, Precisionary Instruments, Natick, MA, USA) and collected in cold PBS. For immunostaining, each slice was placed in PBST (PBS + 1% Triton X-100) with 5% normal horse serum for 3 h and then incubated with primary antibody at 4 °C for 72 h (rabbit anti-C-Fos 1: 2,000, cell signaling; rabbit anti-Tyrosine Hydroxylase (anti-TH) 1: 2,000, Novus; Guinea pig Anti-KCNK2 (TREK-1) 1:200, Alomone Labs, Rabbit anti-GABA (A) α Receptor 1:400, Alomone. Slices then underwent three washes for 20 min each in PBS, followed by 3 h incubation with secondary antibodies (1:500 AlexaFlour488 donkey anti-sheep, (lot#139298) Jackson ImmunoResearch; 1:200 CY5 (for GABA) donkey anti-rabbit, (lot#138569) Jackson ImmunoResearch) and 1:500 CY3 donkey anti-Guinea pig, (lot#134844) Jackson ImmunoResearch). Following another 3 washes with PBS, the brain slices were transferred onto glass slides and mounted with a mounting medium (VECTASHIELD containing DAPI nuclear stain) followed by mounting and coverslipping onto microscope slides. Some clear, colorless nail polish was used to seal the edges of the coverslips.

## Confocal imaging

Fixed slice confocal imaging was performed on a Leica TCS SP8 confocal microscope and Leica DMi 8 at Jacobs School of Medicine and Biomedical Sciences of the University at Buffalo microscopy facility.

## Statistics

To evaluate statistical significance, data were subjected to a Student’s *t*-test or ANOVA, and in the CPP behavior experiments, the investigators were blinded to the different mouse genotypes. Statistical significance was set at P≤0.05. All data are presented as means ± S.E.M. unless noted otherwise.

## AUTHOR CONTRIBUTION

J.L. performed behavioral, biochemical, and imaging experiments, analyzed the data and wrote the paper;

R.M. performed cell culture imaging and partial behavior experiments and analyzed the data. M.H. participated in some behavioral experiments and analyzed the data. S.P performed genotyping experiments and partial imaging experiments. A.P. designed the experiments, supervised the project, and wrote the paper.

## ACKNOWLEDGEMENTS

We thank Dr. Andreas Schwingshackl for sharing the TREK1 KO mice with us. We thank Wade J. Sigurdson for the excellent technical support of confocal imaging from the Jacobs School of Medicine and Biomedical Sciences of the State University of New York at Buffalo. We also thank Denise Ferkey for the support of the Wide-field fluorescence microscope from the Department of Biological Sciences of the State University of New York at Buffalo. This work was supported by the grant from the National Institutes of Health (MH094730) to A.P.

