## Supplemental Figures for "Deep Brain Magnetothermal Silencing of Dopaminergic Neurons via Endogenous TREK1 Channels Abolishes Place Preference in Mice"

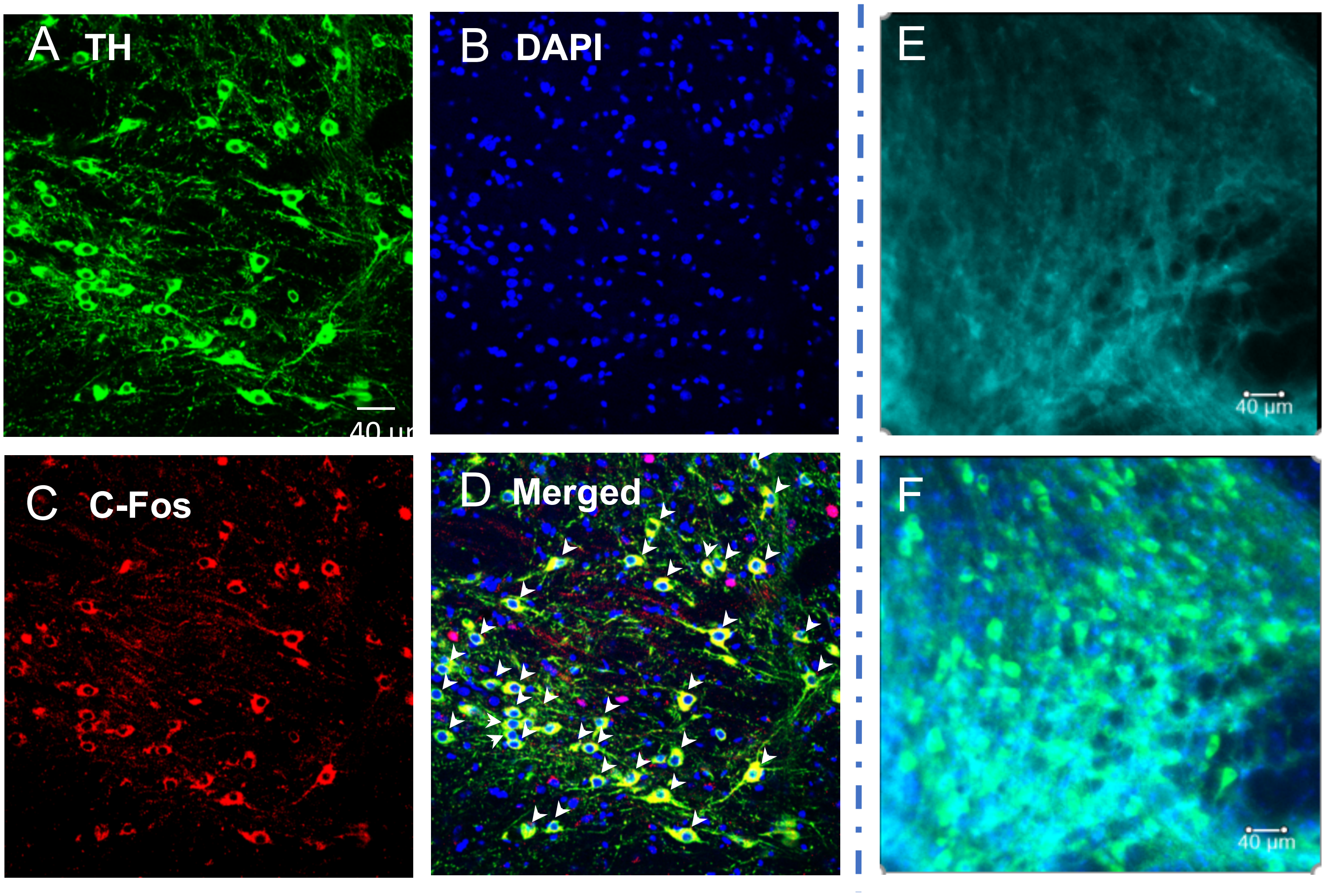


**Figure S1**: Dopamine neurons colocalizated with nuclear, and C-Fos staining in VTA brain slice, and the GABA colocalization with Dopamine neurons. Related to Figure 1. (A) The TH staining for dopamine neurons in the VTA brain region in C57 mice. (B) Nuclear staining with DAPI. (C) C-fos staining on the membrane in the dopamine neurons. (D) The merged image is in the same view in the VTA brain region. (E) GABA staining in the VTA brain region. (F) The merged image of GABA and dopamine neurons in the VTA Brain region

**Figure S2**: Reversible process of neuronal activity during mice freely walk from dark to bright or reverse through a real-time recording by fiber photometry. Related Figure 1. (A) Fluorescence activity signals were recording during mouse walk from dark to bright area. (B) Fluorescence activity signals were recording during of mouse walk from bright area to dark area. (C) Quantification of the VTA neurons' activity for (A) and (B).

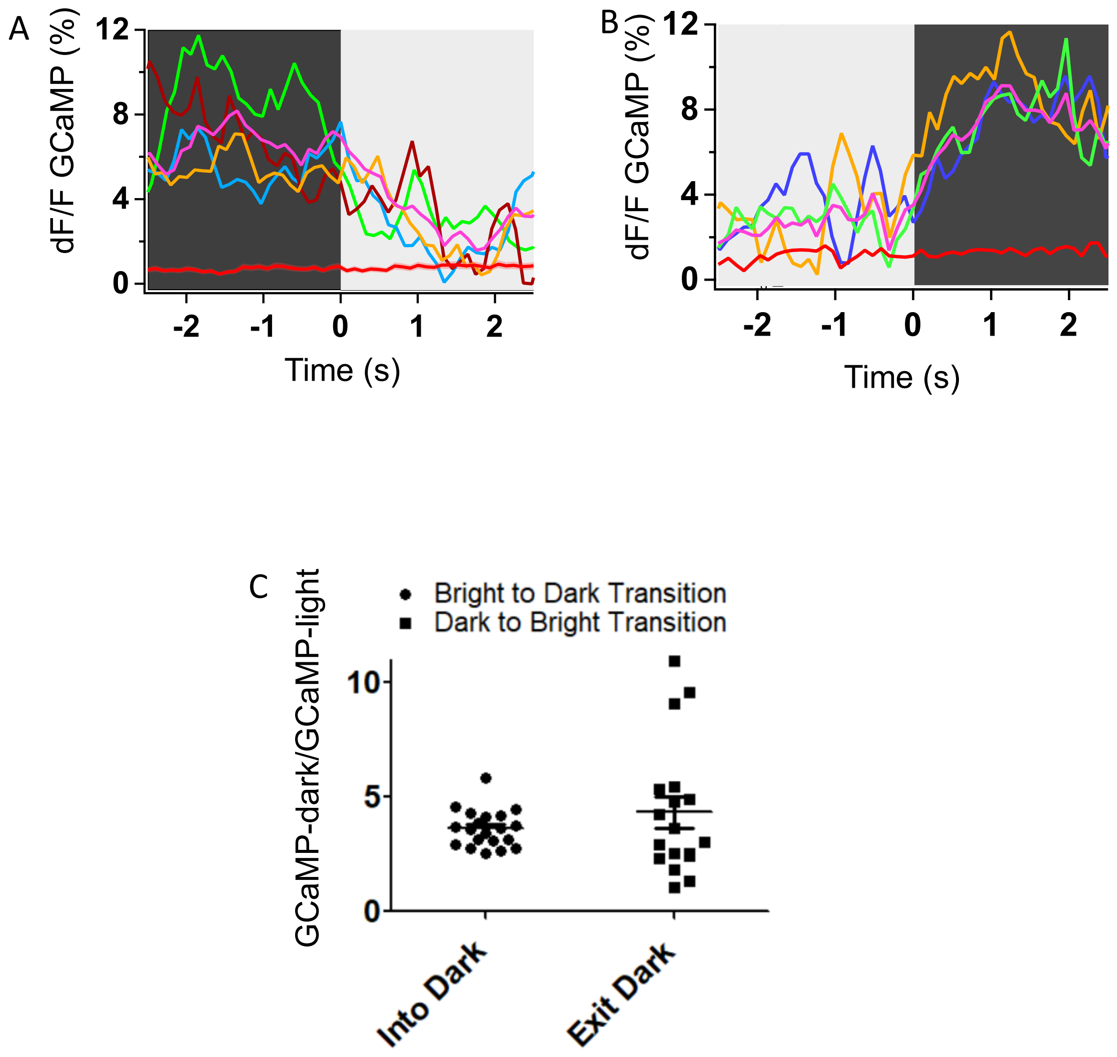
